# CryoVesNet: A Dedicated Framework for Synaptic Vesicle Segmentation in Cryo Electron Tomograms

**DOI:** 10.1101/2024.02.26.582080

**Authors:** Amin Khosrozadeh, Raphaela Seeger, Guillaume Witz, Julika Radecke, Jakob B. Sørensen, Benoît Zuber

## Abstract

Cryo-electron Tomography (Cryo-ET) has the potential to reveal cell structure down to atomic resolution. Nevertheless, cellular cryo-ET data is often highly complex, and visualization, as well as quantification, of subcellular structures require image segmentation. Due to a relatively high level of noise and anisotropic resolution in cryo-ET data, automatic segmentation based on classical computer vision approaches usually does not perform satisfactorily. For this reason, cryo-ET researchers have mostly performed manual segmentation.

Communication between neurons relies on neurotransmitter-filled synaptic vesicle (SV) exocytosis. Recruitment of SVs to the plasma membrane is an important means of regulating exocytosis and is influenced by interactions between SVs. Cryo-ET study of the spatial organization of SVs and of their interconnections allows a better understanding of the mechanisms of exocytosis regulation.

Extremely accurate SV segmentation is a prerequisite to obtaining a faithful representation of SVs state of connectivity. Hundreds to thousands of SVs are present in a typical synapse, and their time-consuming manual segmentation is a bottleneck in this analysis.

Several attempts to automate vesicle segmentation by classical computer vision or machine learning algorithms have not yielded robust results. We addressed this problem by designing a workflow consisting of a U-Net convolutional segmentation network followed by post-processing steps. This combination yields highly accurate results. Furthermore, we provide an interactive tool for accurately segmenting spherical vesicles in a fraction of the time required by available manual segmentation methods. This tool can be used to segment vesicles that were missed by the fully automatic procedure or to quickly segment a handful of vesicles while bypassing the fully automatic procedure. Our pipeline can in principle be used to segment any spherical vesicle in any cell type as well as extracellular vesicles.

## Introduction

The fine architecture of cells can be investigated by cryo-electron tomography (cryo-ET) [1]. Cellular structures are preserved down to the atomic scale through vitrification and observation of the samples in a fully hydrated state. When a macromolecule is present in a sufficient number of copies in the cells imaged by cryo-ET, it is possible to obtain its atomic structure in situ using subtomogram averaging [2,3]. Cellular cryo-ET datasets are usually extremely complex, making them difficult to analyze. This is aggravated by the sensitivity of biological samples to electron radiation, which limits the signal-to-noise ratio in cryo-ET datasets [4]. Tomographic reconstructions are generated from a series of images of the sample acquired at different viewing angles. The geometry of the samples prevents acquisition at certain angles, resulting in anisotropic spatial coverage. The resolution in the directions close to the axis of the electron beam incident on the untilted sample is strongly reduced.

This effect, commonly referred to as the missing-wedge artifact, further complicates data analysis. In particular, organelles fully bounded by a membrane appear to have holes at their top and bottom (relative to the electron beam axis) [4].

The synapse is a specialized cellular contact at which information is transmitted from a neuron to another, the presynaptic and postsynaptic synapses, respectively. In most cases, the signal is transmitted by the release of neurotransmitters into the intercellular space. Neurotransmitters are stored in SVs and are released following the fusion of an SV with the presynaptic plasma membrane. A synapse contains hundreds of SVs and their mobility and recruitability for neurotransmitter release depends on inter-vesicle interactions through so-called connector structures [5,6,7,8]. The characterization of these interactions can be performed automatically with the Pyto software, which implements a hierarchical connectivity approach to segment connectors [9]. For accurate connector segmentation, precise segmentation of SVs is a prerequisite. To date, SV segmentation has been performed manually, but given the large number of SVs per dataset, it is an extremely time-consuming process. Typically, one person spends 3 to 8 working days segmenting a single dataset. Attempts to perform this task automatically based on classical computer vision algorithms have not yielded sufficiently accurate results [10].

To alleviate this situation, we decided to develop an approach based on deep learning. Convolutional neural networks (CNN) have been successfully employed to segment cryo-ET data [11]. Although sufficient for visualization purposes, this approach has not met the requirements to segmenting tethers and connectors in Pyto. Later on, Imbrosci et al. described accurate SV segmentation of transmission electron microscopy images using CNN, but this approach is limited to 2-dimensional (2D) images of resin-embedded synapses [12]. In the former study, cryo-ET data are decomposed in individual 2D slices, which are handed as separate inputs to the CNN. The independent output 2D prediction images are then reassembled in a 3-dimensional (3D) stack [11]. As discussed above, membranes oriented approximately parallel to the plane of the 2D tomographic images are not resolved. In the absence of contextual knowledge of the other 2D images, the CNN fails to segment these regions of the vesicles. Hence, spherical vesicles appear open, whereas we expect closed spherical objects. Recently Zhou et al. addressed this issue by implementing a downstream fitting step based on a Gaussian process approach, allowing for the smooth closure of the membranes [13]. Ideally, 3D networks should be used to segment 3D cryo-ET data. In this context, several groups have published applications of 3D networks in cryo-ET for other tasks, such as particle picking and classification, in order to perform subtomogram averaging [14,15,16]. However, these papers have not focused on accurately segmenting membranes in cryo-ET data.

We opted to employ a 3D U-Net CNN to process 3D images as input [17]. Weigert et al. [18] implemented a U-Net for content-aware restoration (CARE) of 3D fluorescence microscopy datasets. They showed that it can restore information from anisotropic and very noisy datasets. Such networks have been used in the last couple of years in cryo-ET analysis, mainly to perform denoising and object detection [14,15,19]. We implemented a 3D U-Net based on CARE building blocks and trained it with manually segmented datasets. This method provided good accuracy and was not strongly affected by the missing wedge artifact. Nevertheless, it was not sufficient for our downstream Pyto analysis.

Hence, we developed a post-processing method, which transforms the segmented objects into spheres and refines their radius and center location. The workflow includes outlier detection based on the radial profile features of the segmented objects. Then, these mis-segmented vesicles can be either removed or refined. This leads to a substantial improvement in accuracy, which is reflected in Pyto performances comparable to those obtained after manual SV segmentation. We also introduce a semi-automatic method to quickly fix wrongly segmented or missed SVs.

Although our set of procedures was developed with the use case of SV segmentation in mind, it can be used to segment any other types of biological spherical vesicles, such as transport vesicles, secretory vesicles, endocytic vesicles, and extracellular vesicles. Furthermore, with only small modifications it could be extended to extremely accurate segmentation of other membrane-bound organelles or to the plasma membrane.

## Results

### Overview of training and work**fl**ow

In view of the effort required for the manual segmentation of SVs, we decided to develop an automatic segmentation procedure. Since we had previously manually segmented a number of tomograms with the program IMOD, we could use these segmentations as the ground truth [20]. We trained a U-Net with a set of 9 segmented tomograms of rat synaptosomes (see Materials and Methods).

We sought to further improve segmentation accuracy by feeding the probability mask output by the U-Net to a series of post-processing steps (Figure 1). Three sets of tomograms were used to assess the performances of the pipeline:

**Figure 1:**
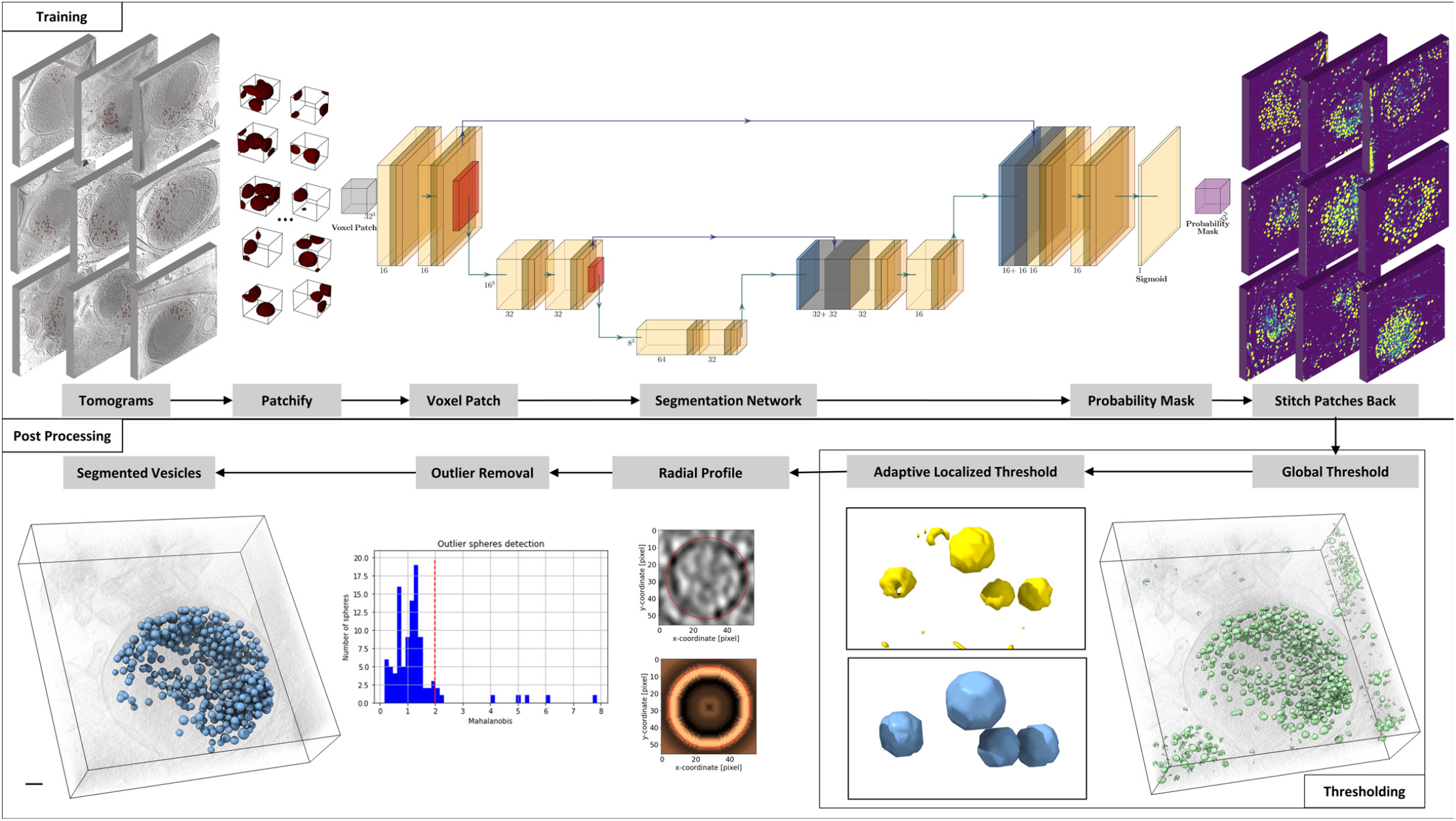
Pipeline of automatic segmentation. a) Tomograms b) Splitting in 3D patches c) Segmentation Network/ trained U-Net d) probability masks e) stitching patches back together f) thresholding h) radial profile i) outlier removal j) Segmented Vesicles

1. Train tomograms : The 9 rat synaptosome tomograms that have been used for U-Net training
2. Test tomograms: 9 additional rat synaptosome tomograms
3. Generalization test tomograms: 12 mouse primary neuronal culture tomograms.

Each tomogram was split into patches of 32^3^ voxels. These patches were fed in the U-Net, which outputs a probability mask for those patches. To obtain a complete probability mask, the patches are stitched back together (Figure 1, Figure EV1). The probability mask was then binarized with a global threshold step.

We noticed that some vesicles were not segmented accurately. Indeed, some vesicles that were in close proximity were misidentified as a single entity. Separating them necessitated adjusting the detection threshold to a more stringent value. (see “Adaptative local thresholding” section in Materials and Methods). Additionally, there were instances where the detection captured only a fraction of a vesicle, which required loosening the threshold for a more accurate segmentation. Yet the assignment of the correct label to each segmented vesicle was essential for the next steps.

### Sphericalization and radial pro**fi**le-based re**fi**nement

Although at first glance the segmentation looked good after these steps, we noticed that it was not extremely accurate. For example, the vesicles were not always centered in the segment or the radius of the segment was inexact. Very often, the vesicle segment looked shrunk in the z-direction, whereas the actual vesicles were spherical. This would be highly problematic for automatic connector and tether segmentation. To address these issues, we represented each vesicle as a sphere. We determined the center and radius of the sphere as described in the Materials and Methods section.

We then performed a radial averaging of the intensity around the center of the sphere. And we adjusted iteratively the position and radius of the sphere to match the actual structure in the tomogram (Figure 2). The radial profile refinement is a pivotal tool as it ensures that the segmented vesicles are a true representation of their form in the tomogram.

**Figure 2:**
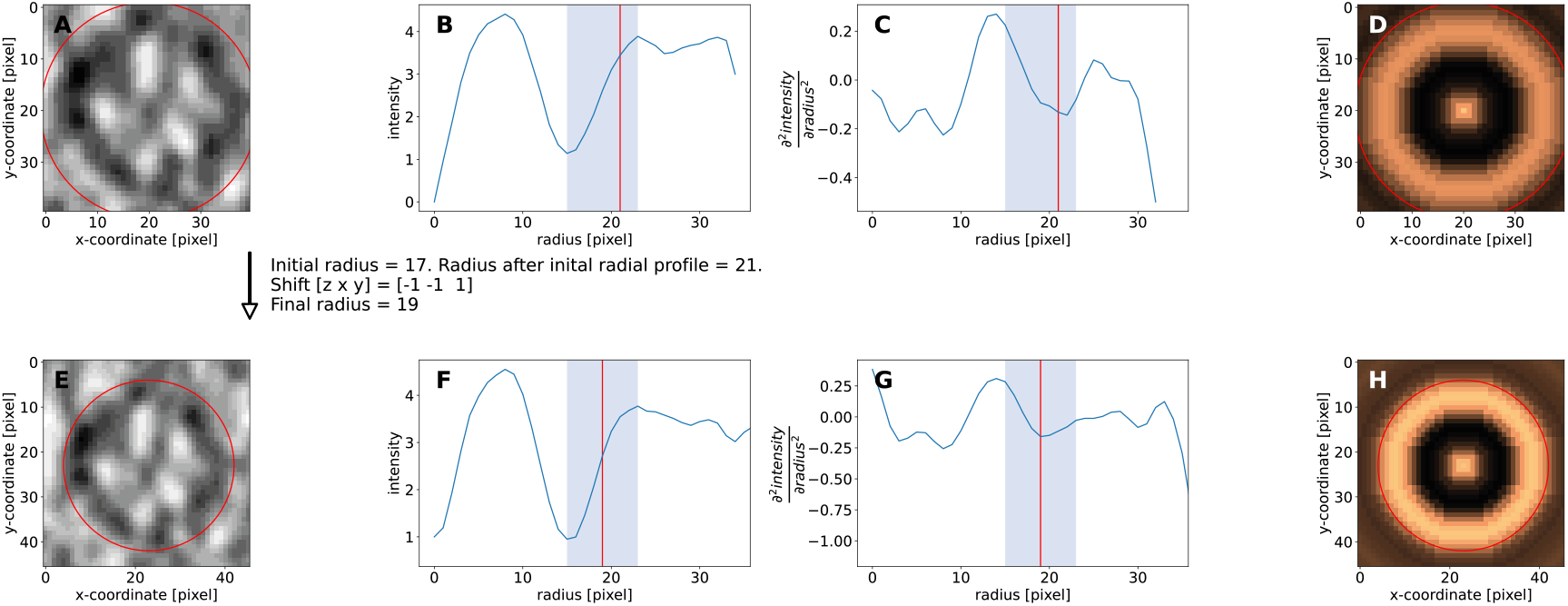
Vesicle radius and position refinment through radial profile and cross-correlation. (A) Initial segmentation of a vesicle. (B) Radial Profile. Blue range is from membrane center to outer white halo center. This is defined as the search range for the optimal radius. (C) second derivative of radial profile, used to define the exact edge of the membrane. (D) Central cross-section in the three-dimensional radial average of the vesicle in its initial position. (E-H) Same as (A-D) after refinement.

### Outlier detection and re**fi**nement

Despite the improvement brought by the radial profile refinement, some vesicles were still not segmented accurately. By quantifying several parameters of the segmented vesicles, such as radius, membrane thickness, membrane intensity, lumen intensity, we were able to spot outliers using multivariate statistics (see Materials and Methods). An example of outlier detection is shown in Figure 3. In this example, three outliers are highlighted. Outlier 1 (red, top row) corresponds to the mistaken segmentation of a non-vesicular membrane-bound structure. The high mahalanobis distance of this outlier can be explained by a vesicle radius, membrane thickness, and intensity that are very different from the average of the dataset. Outlier 2 (green, middle row) is correctly segmented but is flagged for its abnormally high radius. Indeed, both the membrane thickness and intensity are close to the average of the dataset but the highly increased radius leads to a high mahalanobis distance. Outlier 3 (blue, bottom row) is initially detected but is misplaced. Its radius was not divergent from the average but its membrane thickness and intensity were. We could then refine these outliers or remove them if refinement failed.

**Figure 3:**
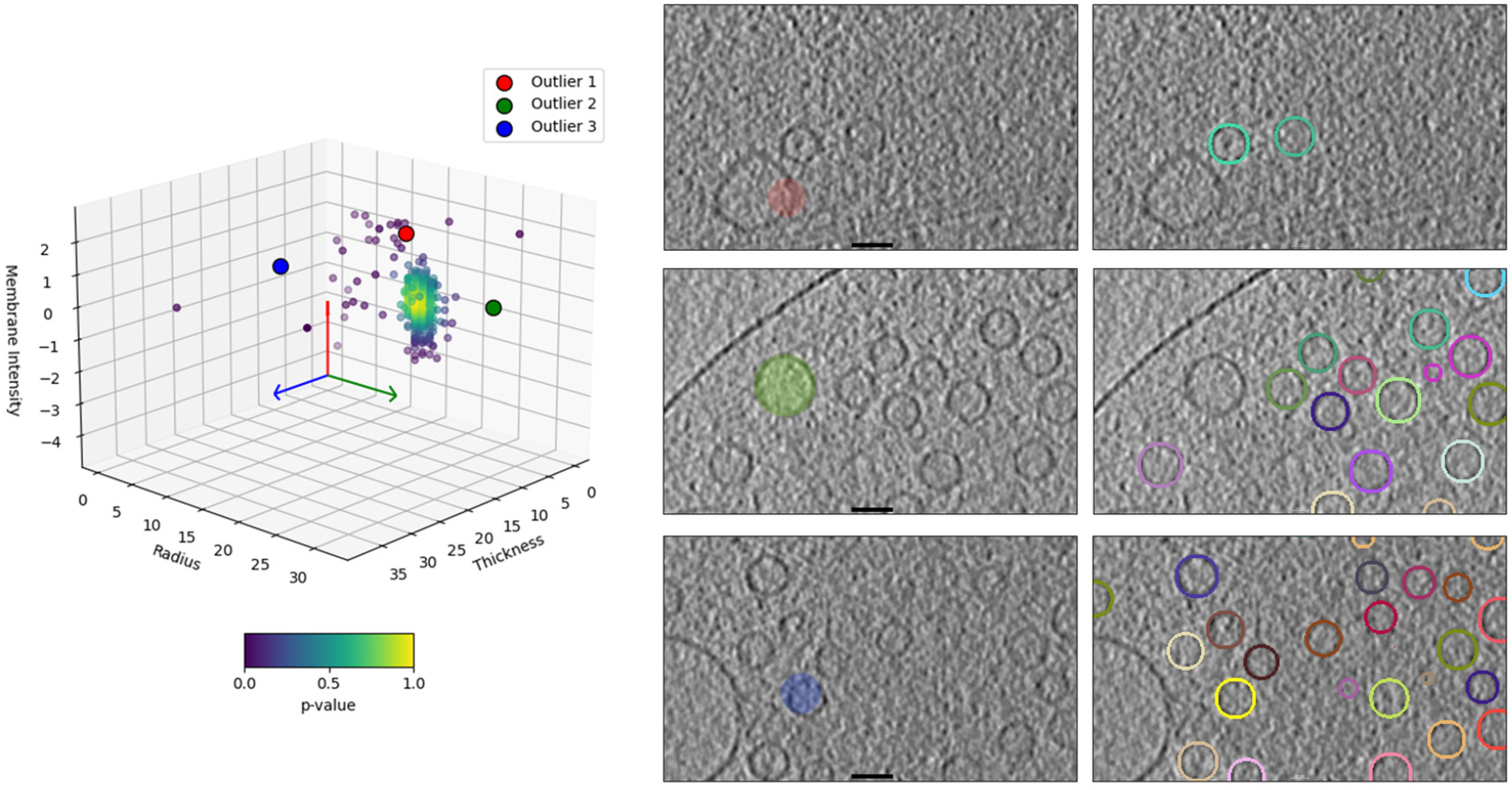
Multidimensional Outlier Detection. The scatter plot (left panel) represents vesicle features in the space defined by membrane intensity, radius, and thickness, with points colored according to the p-value of their Mahalanobis distance, identifying potential outliers. Central panels: outliers are highlighted. Right panels: outliers have been either removed (top and middle row) or fixed by refinement(bottom row). In addition the right panels show the final vesicle segmentation boundaries. Bars, 100 nm.

### Performance and generalization

The performance of all steps was quantitatively assessed by comparing the obtained segmentation with the ground truth using the Dice coefficient metric (see Materials and Methods). The Dice coefficient of the probability mask was 0.64±0.11 for the train tomograms, 0.75±0.06 for the test tomograms, and 0.69±0.09 for the generalization test tomograms (Table 1, Table 2, Table 3, Table EV1, Table EV2, and Figure 4). The probability mask was then binarized with a global threshold step, which led to a Dice of 0.78±0.04 and 0.80±0.04 in the train and test datasets, respectively, while it led to a slight decrease dice of 0.66±0.09 in the generalization test dataset. You do not see instantly that localized thresholding step affect the performance significantly however, it is necessarily to avoid false negative vesicles. The Radial profile refinement and final outlier removal steps led to a Dice coefficients of 0.86±0.05, 0.83±0.05, and 0.79±0.09, respectively.

**Table 1:** Evaluation of the segmentation on the synaptosomal train set. Mask Dice: Mask Dice coefficient for the predicted mask; Final Label Dice: Dice coefficient after post-processing; δ d: average relative diameter deviation over all correctly detected vesicles; Δ c: center residual error average and standard deviation (nm); Number of Vesicles: number of expected vesicles; TP: True Positive; FN: False Negative; FP: False Positive; F1-score: F1-score based on TP, FN, and FP.

| Dataset | Mask Dice | Final Label Dice | $\delta_d$ | $\Delta_c(nm)$ | Number of Vesicles | TP | FN | FP | F1-score |
| --- | --- | --- | --- | --- | --- | --- | --- | --- | --- |
| SC 1 | 0.44 | 0.73 | 0.07 | $2.55 \pm 1.56$ | 223 | 198 | 26 | 49 | 0.841 |
| SC 2 | 0.80 | 0.90 | 0.05 | $2.12 \pm 1.06$ | 105 | 103 | 2 | 1 | 0.986 |
| SC 3 | 0.67 | 0.90 | 0.05 | $1.86 \pm 1.24$ | 128 | 127 | 1 | 6 | 0.973 |
| SC 4 | 0.62 | 0.89 | 0.03 | 1.78±0.92 | 144 | 141 | 3 | 4 | 0.976 |
| SC 5 | 0.58 | 0.87 | 0.04 | 1.86±1.00 | 214 | 209 | 5 | 13 | 0.959 |
| SC 6 | 0.56 | 0.84 | 0.04 | 1.92±1.05 | 104 | 102 | 2 | 16 | 0.919 |
| SC 7 | 0.78 | 0.88 | 0.06 | 1.86±0.90 | 184 | 184 | 0 | 16 | 0.958 |
| SC 8 | 0.75 | 0.90 | 0.05 | 1.70±0.93 | 132 | 126 | 6 | 1 | 0.973 |
| SC 9 | 0.59 | 0.87 | 0.05 | 1.87±0.91 | 135 | 132 | 3 | 14 | 0.940 |
| **Average** | **0.64±0.11** | **0.86±0.05** | **0.05** | **1.95±1.08** | **152.22** | **97.00%** | **3.00%** | **7.30%** | **0.95±0.04** |

**Table 2:** Evaluation of the segmentation on the synaptosomal test set. (same sample type as the train set). For the meaning of the columns, see Table 1.

| Dataset | Mask Dice | Final Label Dice | $\delta_d$ | $\Delta_c(nm)$ | Number of Vesicles | TP | FN | FP | F1-score |
| --- | --- | --- | --- | --- | --- | --- | --- | --- | --- |
| SC 10 | 0.75 | 0.88 | 0.07 | 1.86±1.18 | 129 | 123 | 6 | 5 | 0.957 |
| ST 1 | 0.75 | 0.83 | 0.11 | 2.66±1.52 | 699 | 687 | 12 | 33 | 0.968 |
| ST 2 | 0.74 | 0.77 | 0.11 | 2.27±1.84 | 122 | 117 | 5 | 2 | 0.971 |
| ST 3 | 0.72 | 0.74 | 0.11 | 3.64±2.22 | 434 | 397 | 37 | 57 | 0.894 |
| ST 5 | 0.77 | 0.85 | 0.08 | 2.20±1.26 | 535 | 526 | 9 | 25 | 0.969 |
| ST 6 | 0.60 | 0.83 | 0.07 | 2.02±1.12 | 373 | 353 | 20 | 42 | 0.919 |
| ST 7 | 0.80 | 0.83 | 0.06 | 2.22±1.14 | 110 | 107 | 3 | 9 | 0.947 |
| ST 8 | 0.83 | 0.91 | 0.04 | 2.09±1.04 | 100 | 99 | 1 | 2 | 0.985 |
| ST 10 | 0.77 | 0.86 | 0.05 | 1.96±1.04 | 77 | 74 | 3 | 6 | 0.943 |
| **Average** | **0.75±0.06** | **0.83±0.05** | **0.08** | **2.32±1.43** | **286.56** | **96.30%** | **3.70%** | **6.10%** | **0.95±0.03** |

**Table 3:**
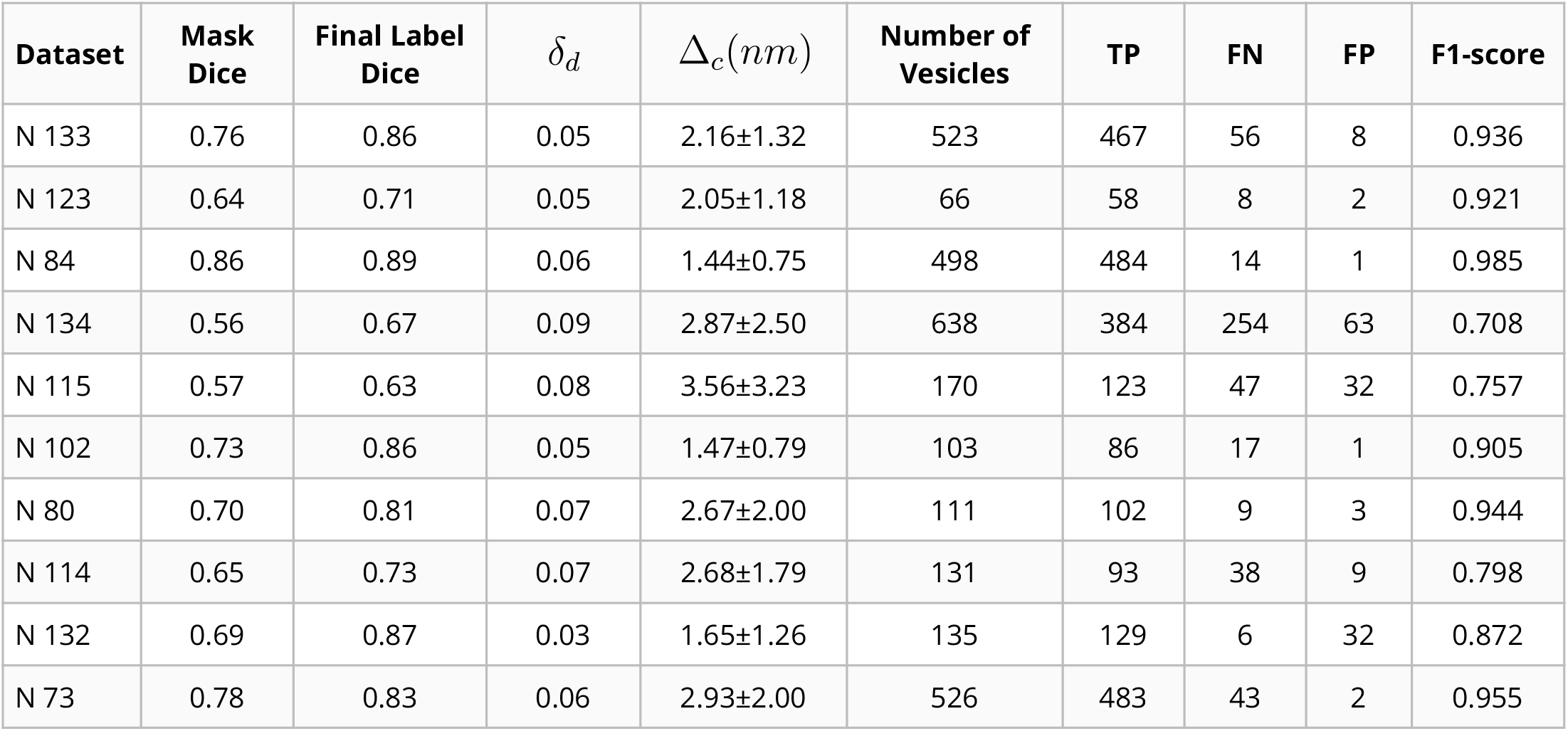
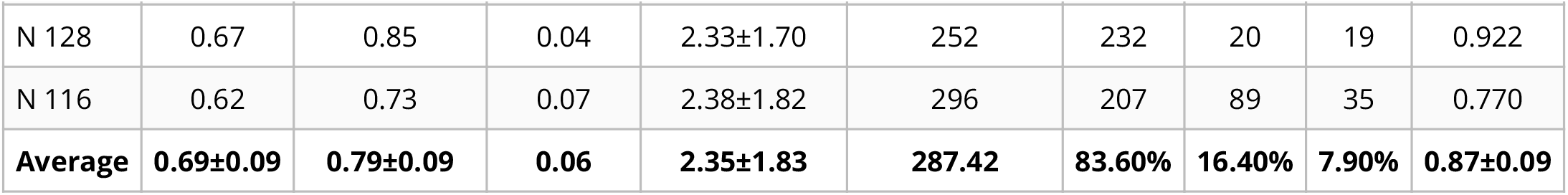
Evaluation of the segmentation on the neuronal generalization test set. (different sample type as the train set). For the meaning of the columns, see Table 1.

**Figure 4:**
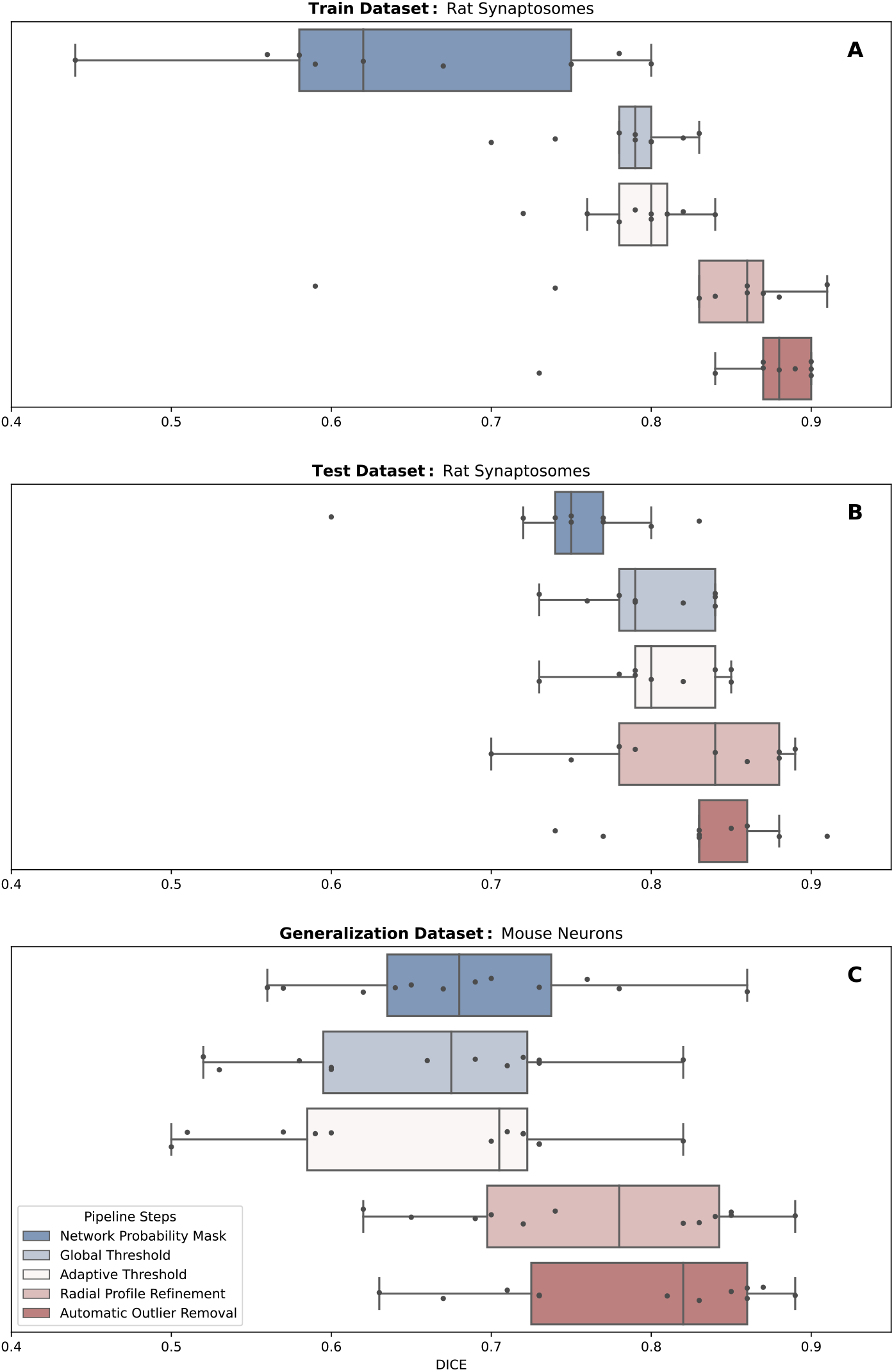
Dice development during post-processing. Dice development at different post-processing steps of initially predicted mask (different colors correspond to different tomograms): A) synaptosomal train set B) synaptosomal test set c) neuronal generalization test set

In addition to the Dice metric, which is a voxel-wise evaluation, we performed a vesicle-wise evaluation. Namely, we quantified vesicle diameter deviation, and center residual (Figure 4, Table 1, Table 2, and Table 3). Results show that our method transfers well across datasets even without fine-tuning which shows robustness and generalization.

### Downstream analysis and application

Traditional manual segmentation, while precise, is time-consuming and often limited in scope. In previous cryo-ET studies of presynaptic terminals, the analysis of spatial organization was restricted within 250 nm of the active zone in order to keep segmentation time reasonable. This limitation inherently narrows the scope of synaptic analyses. The advent of deep learning-based segmentation offers a promising alternative, providing both speed and scalability. We were able to segment full synapses in a fraction of the time that it manual segmentation would take. Furthermore, we have implemented an interactive tool with Napari [21] that enables to swiftly rectify false positives and negatives by adding or removing vesicles as needed, ensuring the highest level of accuracy in the final segmented output. A comparison of manual and automatic segmentation is shown in Figure 5. Pyto is a software package designed for the analysis of pleomorphic membrane-bound molecular complexes in 3D images, particularly in the context of synaptic cryo-ET. A key feature of Pyto is its ability to accurately segment connectors and tethers within the pre-synaptic terminal, a task that requires a high level of vesicle segmentation precision. This segmentation process is hierarchical and connectivity-based, detecting densities interconnecting vesicles (connectors) and densities connecting vesicles to the active-zone plasma membrane (tethers). CryoVesNet has been designed to be compatible with Pyto and an application is demonstrated in Figure 6. This enables us to extract a wealth of structural information to better understand the structural basis of synaptic vesicle exocytosis regulation.

**Figure 5:**
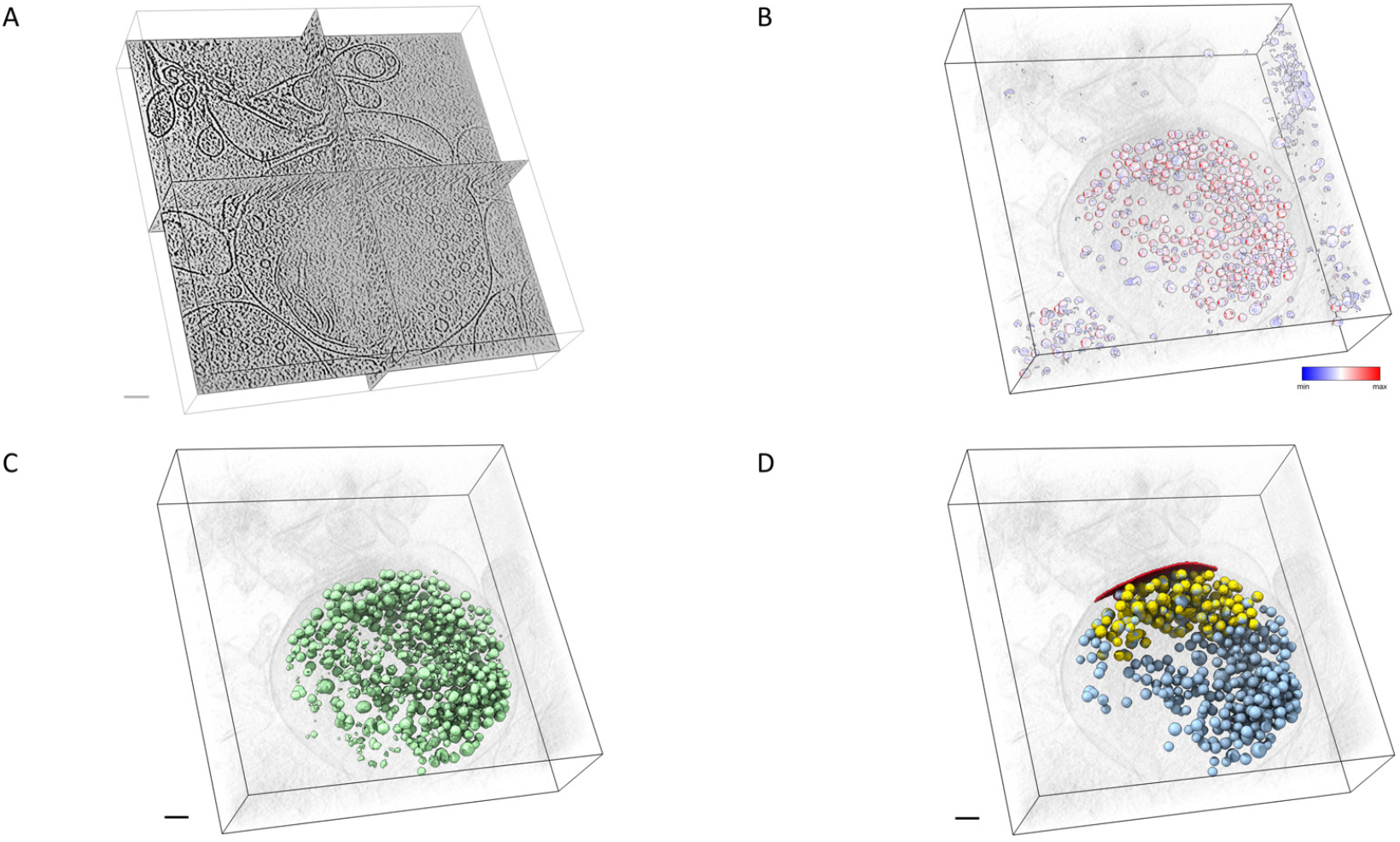
Comparison of manual and automatic vesicle segmentation. A: 3D representation of a synaptasome density map, visualized in an orthogonal view.B: Probability map output by the neural network. The map is represented with a color gradient ranging from blue (0.9968) to red (1), indicating the likelihood of synaptic vesicle presence. C: Segmentation after global threshold optimization. D: The final representation of synaptic vesicle segmentation post-processing. CryoVesNet segmented vesicles are depicted in light blue, combined with expert annotation in yellow (restricted within 250 nm of the active zone shown in red). Bar, 100 nm.

**Figure 6:**
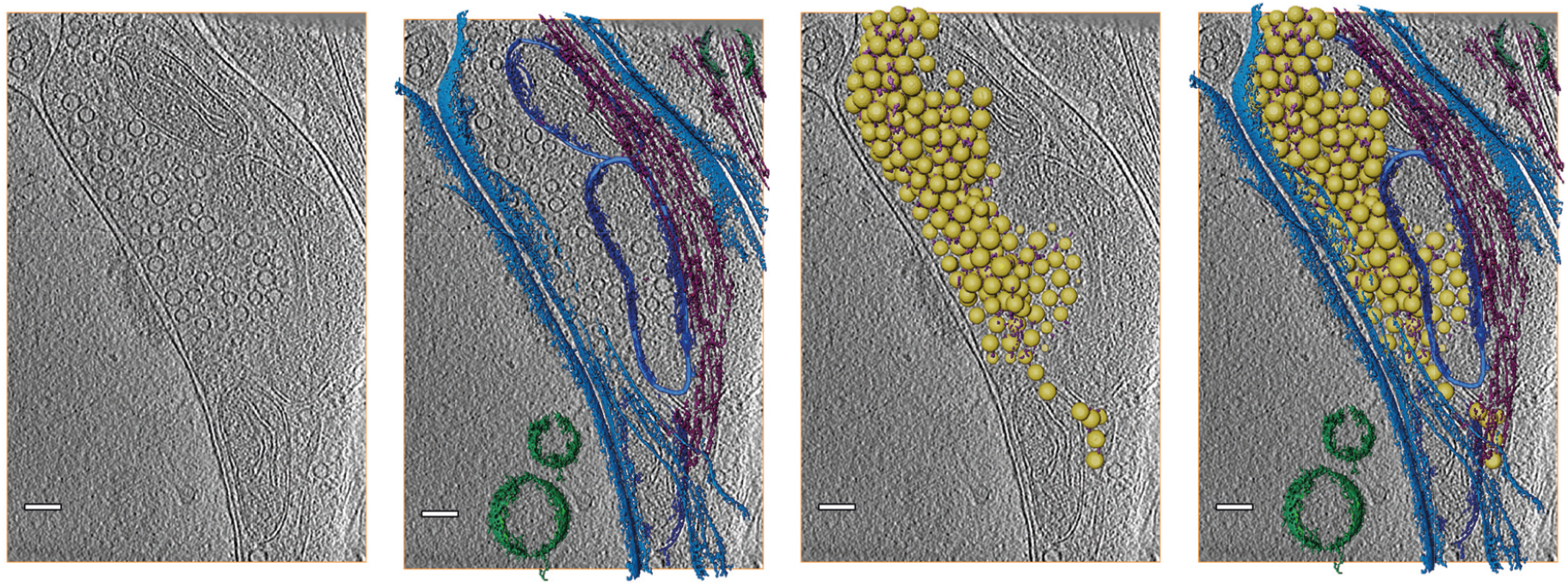
3D model of a cultured mouse neuron synapse. A: slice through a tomogram. B: Segmentation of plasma membrane (light blue), mitochondria (dark blue), endosomes (green), microtubules (dark magenta). C: Vesicles (yellow) and connectors (pink) segmented with CryoVesNet and Pyto, respectively. D: Combination of B and C. Bar, 100 nm.

## Discussion

### Synaptic Vesicles: Molecular Interactions and Functions

Synaptic vesicles (SVs) play a central role in neurotransmission, facilitating the release of neurotransmitters into the synaptic cleft. These vesicles undergo a series of molecular interactions with various protein complexes, transitioning from a tethered to a primed state, and eventually to neurotransmitter release through exocytosis. Synapsins have been identified as key proteins in regulating the availability of SVs for exocytosis. It has been hypothesized that synapsins cross-link SVs, thereby preventing their premature release.

Cryo-ET emerges as a powerful tool to address these challenges, offering unparalleled insights into the molecular architecture of synapses. Accurate segmentation of structures such as vesicles, connectors, and tethers is essential for a comprehensive understanding of synaptic function. Cryo-ET, however, is not without its challenges. The technique suffers from a high level of noise and anisotropic resolution (known as the missing wedge phenomenon), which complicates data analysis and interpretation. Addressing these challenges is crucial for obtaining clear and accurate tomographic reconstructions.

### CryoVesNet: Automatic Vesicle Segmentation in Cryo-ET

By utilizing a U-Net architecture trained on manually segmented tomograms and postprocessing steps, we have developed a system that can efficiently and accurately segment synaptic vesicles in tomographic datasets. In particular, CryoVesNet is uniquely insensitive to the missing wedge and can segment complete vesicles even if the membrane is not fully visible in the tomogram. The results obtained from our method, as evidenced by the Dice coefficient and other evaluation metrics, demonstrate its robustness and accuracy. Notably, our method ability to generalize across different datasets, namely from rat synaptosomes and to primary neuronal cultures, underscores its versatility and potential for widespread application. It is not restricted to synaptic vesicles but can be applied to any spherical membrane-bound organelle. It is interesting to note that the Dice coefficient of the U-Net output (probability mask) was better for the test set and generalization set than the train set. Consequently the train set benefited more strongly from the post-processing steps than the test and generalization sets in term of Dice coefficient.

The use of both global and adaptive localized thresholding techniques further refines the segmentation, addressing challenges posed by closely packed vesicles. Our results highlight the effectiveness of our post-processing steps, including radial profile refinement and the removal of outliers. The radial profile, in particular, ensures that the segmented vesicles closely match their actual structure in the tomogram, providing a more accurate representation. Furthermore, our method compatibility with software like Pyto, which is designed for the analysis of pleomorphic membrane-bound molecular complexes in 3D images, enhances its utility. By integrating our segmentation approach with tools like Pyto, researchers can gain deeper insights into vesicle interactions. Significantly, with minimal adjustments, our method could be adapted to achieve highly precise segmentation of any membrane-enclosed organelle or the plasma membrane itself.

## Conclusion

In conclusion, CryoVesNet for automatic segmentation in cryo-ET represents a significant step forward in the study of synaptic vesicles and their associated structures. By combining the power of deep learning with optimized post-processing techniques, we offer a solution that is both efficient and precise. As the field of structural cell biology continues to evolve, tools like ours will play a crucial role in advancing our understanding of complex cellular structures and processes.

## Materials and methods

### Cryo-electron tomography datasets

In this study, we used datasets originating from either rat synaptosomes or mouse primary neuron cultures. They represent a total of 30 tomograms with heterogeneous pixel sizes, defocus and resolution and we split them in three groups: 1. Train set: 9 synaptosome tomograms were used for training. 2. Test set: 9 independent synaptosome tomograms were used for testing. 3. Generalization test set: 12 neuron tomograms were used for assessing transfer learning potential. The preparation procedure of the samples from which the datasets were obtained as well as the biological analysis of these datasets was previously reported [22].

### Manual segmentation and automatic interboundary segment detection

Manual segmentation of SVs, the presynaptic cytoplasm, and the active zone plasma membrane was done in IMOD [20]. SVs were segmented as spheres. The presynaptic cytoplasm marked the region to be analyzed by Pyto [9]. Later on, we refer to this region as the cytoplasmic segmentation region. It consisted of the volume comprising the active zone and the cluster of SVs. The analysis by Pyto was essentially the same as described previously [5,9]. In short, the segmented region is divided in 1 voxel thick layers parallel to the active zone for distance calculations. A hierarchical connectivity segmentation detects densities interconnecting boundaries. The boundaries were synaptic vesicles and the active zone plasma membrane. Detected intervesicular segments are termed connectors and segments connecting vesicles to the active zone plasma membrane are called tethers. Distance calculations respective to SVs were done from SV center. The segmentation procedure is conservative and tends to miss some tethers and connectors because of noise. Consequently, the numbers of tethers and connectors should not be considered as absolute values, but rather to compare experimental groups. As it was done before, an upper limit was set between 2100 and 3200 nm^3^ on segment volume. The tomograms that were used for this analysis were binned by a factor of 2 to 3, resulting in voxel sizes between 2.1 and 2.4 nm.

### Train and validation set generation

In the preparation of our train set, we utilized segmented 3D image volumes. The primary volume was systematically divided into 32^3^ cubic sub-volumes. To ensure the relevance and richness of the data, only those sub-volumes that were sufficiently occupied by vesicles, specifically containing more than 1000 voxels, were retained. 860 sub-volumes were used for training and 100 sub-volumes were used for validation.

### Network architecture and training procedure

We used a U-Net with two downsampling stages and two convolutional layers per stages, with a kernel size of 3, and ReLU activation function based on the open-source CARE framework (Figure 1) [18]. We used the Adam optimizer on a binary cross-entropy loss function. The learning progress was tracked by calculating the Dice coefficient and the loss value after each training epoch (Figure EV2). The Dice coefficient for the train set was initially ∼0.25 and rose to over 0.9 after approximately 50 epochs, while for the validation set, it increased to 0.8. The loss for the train set went from 0.55 and to values below 0.05 after 50 epochs while for the validation set it went from 1 and to slightly below 0.3 after the initial 50 epochs and then rose slightly. The training was done for 200 epochs.

### Probability map construction

Our U-Net model, trained on 32^3^-voxel patches, utilizes a 24-voxel region of interest (ROI). To mitigate tiling effects during testing, the network input can be expanded to accommodate larger volume specifically in this case 64^3^ voxels. The tomogram undergoes padding to align with the ROI, ensuring reduced edge artifacts. Segmentation is executed in tiles, where the U-Net predicts the synaptic vesicle probability for each tile. Only the central part of the segmented patch, corresponding to the ROI, is retained. Finally, the segmented tiles are reassembled, yielding a continuous synaptic vesicle probability map of the entire volume.

### Global thresholding

Segmenting implies turning the probability map into a binary mask. In order to find the optimal threshold value, we iterated through potential threshold values ranging from 0.8 to 1 in increments of 0.01. A binary mask was generated for each threshold. Subsequently, an erosion operation was applied to the binary mask, and the difference between the original and eroded masks produced the vesicle shell. The voxel intensity values of the original image corresponding to this shell were recorded for each threshold. We minimized the average intensity of the shell voxels to determine the optimal threshold value, since the shell of correctly segmented vesicles corresponds to the vesicle membrane, which in cryo-ET appears darker, i.e., with lower intensity values.

### Adaptative local thresholding

Each individual segment was given a label using the scikit-image label method [23]. A majority of vesicles were correctly segmented but we noticed some segments included more than one vesicle. We therefore evaluated each segment with two criteria based on the fact that synaptic vesicles have a homogenous size and are spherical. Firstly we calculated the volume z-score *z* for each segment:

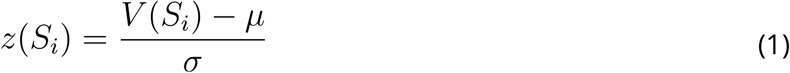

where *S*_*i*_ is the segment *i,V* (*S* _*i*_) is the volume of *S* _*i*_, *μ* the average volume of all segments, and σ the standard deviation of the segment volumes. Secondly, we computed the segment extent *e* as:

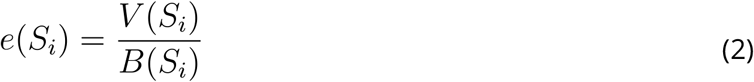

where *B*(*S* _*i*_) is the volume of segment *i* bounding box. The extent of a sphere equals 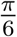. Segments with both a z-score *z*>1 and an extent *e<0*.*25* were considered as potentially comprising more than one vesicle. For each of these segments, the threshold was increased stepwise until two distinct segments were generated. Subsequently, the extent and volume of all segments was evaluated again. Any segment with *e<0*.*25*,or *e<0*.*75*,*Vk* or was discarded. *k* was defined as the volume of a sphere with a radius of 12 nm. This ensured that segments deviating highly from spherical shape and segments with a volume smaller than an acceptable volume *k* were removed.

### Segmentation refinement using radial profile

Even if most synaptic vesicles were detected and well segmented, segmentation accuracy was not sufficient for our downstream application. To improve accuracy, each segment was converted to a spherical segment and its radius and position was refined. Initial spherical conversion was done by setting the center of the sphere *C* at the position of the centroid of the segment, while the radius was defined as half the length of the bounding box longest edge. The segment position and radius were iteratively refined as follows.

1. The radial average ⟨*I*(*d*)⟩ was computed:

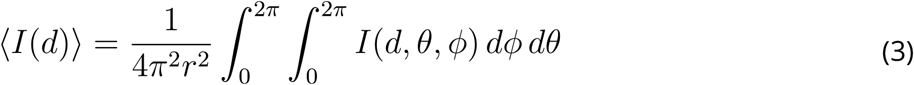

where *d* is the radial distance from the segment center, θ the polar angle, and *ϕ* the azimuthal angle.
2. The radius of the vesicle was updated as:

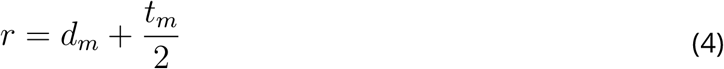

where *d*_*m*_ is the radial distance of center of the vesicle membrane, and *t*_*m*_ the thickness of the vesicle membrane. *d*_*m*_ was defined as the radial distance for which the radial average was minimal. 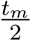 was calculated as the distance between the center of the vesicle membrane and the minimum of the second derivative of the radial profile in the interval between the center of the vesicle membrane and the maximum of the Fresnel fringe outside the membrane.
3. The radial average was back projected in 3-dimension:

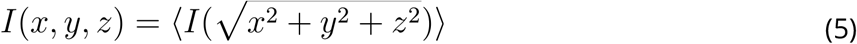

where (*x,y,z*)= (0,0,0) is the coordinate of the segment center.
4. We computed by cross-correlation the shift between the obtained 3-D average and the 3-D image in the cubic box with central coordinates *C* and edge length *l =2r +c*, where *c* is a constant.*C* was updated by subtraction of the shift.
5. Steps 1 to 4 were repeated for a maximum of 10 iterations until convergence or until a total shift of 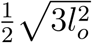, where *l*_*0*_ is the edge length of the initial box. The feature space of predicted vesicle labels was computed, containing membrane thickness *t*_*m*_, membrane intensity *ρ*, and vesicle radius *r. ρ* was defined as the mean intensity of the radial average within the radial distance interval 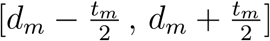.

### Outlier detection and re**fi**nement

Following radial profile calculation, key features, namely thickness, radius, and membrane intensity, were extracted. Using these criteria, the Mahalanobis distance *D*^2^ was calculated for each data point to quantify its distance from the distribution of these features as follows, using Scipy’s implementation [24]:

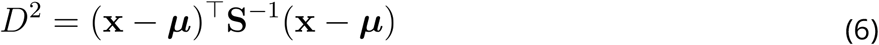

where ***x*** is the vector of the three features, ***μ*** is the mean vector of the features, and ***S***^−1^ is the inverse covariance matrix of the features. The significance of the Mahalanobis distance was interpreted as follows:

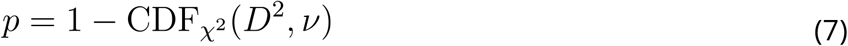

where 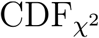 is the cumulative distribution function of the chi-squared distribution, and *v* is the number of degrees of freedom, which is equal to the number of features - 1. The resulting p-value provides a measure of the statistical significance of each data point’s distance (Figure 3).

With the computed p-values, outlier detection and refinement were conducted. Each vesicle with a p-value lower than a given threshold was defined as an outlier. In this study, we empirically set the threshold to 0.3, but other values can be used, depending on the use case. The radial profile and p-value of the outliers was recalculated using a different box size. We performed this step iteratively. At each iteration the box size was made larger by 2x2x2 voxels. For each outlier, the iteration stopped when its p-value was higher than the threshold. A maximum of 10 iterations was performed. Vesicles that did not meet the p-value criteria were removed from the dataset.

### Evaluation Metrics

The evaluation framework was designed to assess the capabilities of the proposed toolbox for automatic synaptic vesicle segmentation. We defined as ground truth the manual segmentation of synaptic vesicles. Evaluation was peformed within the cytoplasmic segmentation region (see “Manual segmentation and automatic interboundary segment detection”). We performed per vesicle evaluation and voxel-wise evaluation. For the former, we defined a vesicle as correctly segmented if the center of the predicted vesicle was located inside the ground truth vesicle. Based on that we calculated an F1 score. For the voxel-wise evaluation, we calculated the Dice coefficient between prediction and the ground truth.

### Voxel-wise evaluation

During training a Dice coefficient for probabilistic subvolume maps was calculated after each epoch as a performance quantification EV2. After reconstructing the probablility map after training or prediction, we employed the Dice metric for the whole tomogram to evaluate the similarity of the predicted probability mask with ground truth 1,2, and 3. The Dice coefficient is defined as:

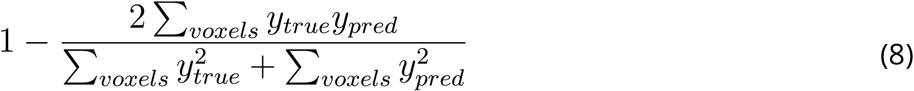

where *y*_*true*_ and *y*_*pred*_ are the ground truth and predicted probability values, respectively, for each voxel.

The Dice coefficient was also employed to monitor all stages of post-processing of the labels, to observe the effect of each post-processing step.

### Vesicle diameter and position deviation

In addition to the F1-score, we also evaluated the precision of our segmentation by calculating the deviation of the estimated vesicle diameter and position from the ground truth. Given the diameter of a ground-truth manually segmented vesicles *d*_*GTi*_ and the predicted diameters of the same vesicles *d*_*pi*_, the average deviation in diameter estimation across all vesicles can be expressed as *δ*_*d*_, where *δ*_*d*_ is calculated as:

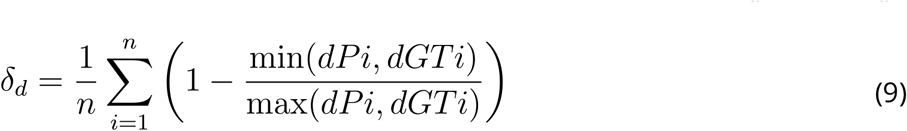

Here, n represents the total number of vesicles, is *dPi* the diameter of i-th vesicle predicted by the segmentation, and *dGTi* is the diameter of the i-th vesicle in the ground truth. This formula offers an insight into the congruence between our estimated diameter and the manual segmentation, with a diminished value of *δ*_*d*_ signifying a closer approximation.

Similarly, the average deviation in position estimation across all vesicles can be expressed as Δ ^*c*^. It corresponds to the average euclidean distance between the center of ground truth vesicles and the center of predicted vesicles.

### Statistical comparison

Multiple pairwise ANOVA comparisons with Benjamini-Hochberg correction were performed on the Dice values summarized in EV1 to assess the statistical significance of the differences between the Dice values [25]. We performed Benjamini-Hochberg correction with the multipletests function implemented in the Python module statsmodels [26]. A list of P-values resulting form pairwise comparisons was input, and multipletests output a list of corrected P-values. The used implementation of the Benjamini-Hochberg correction does not require a false discovery rate to be input. This variation of the original Benjamini-Hochberg correction algorithm was proposed by Yekutieli and Benjamini [27]. If a corrected P-value is smaller than the defined acceptable false discovery rate, then the null hypothesis is rejected, i.e. the difference is considered statistically significant. This algorithm enables to test multiple false discovery rates in one step and its conclusions are exactly the same as the original Benjamini-Hochberg correction algorithm run multiple times with different false discovery rates.

### Computational Setup

All experiments were conducted using 4 x NVIDIA 2080 Ti GPUs with CUDA 10.1. The software environment was set up with Python 3. Key libraries and packages used include TensorFlow 2.4.1 with GPU support and Keras 2.4.3. Image visualization was achieved with UCSF ChimeraX [28] and Amira 2022.2 (Thermofisher Scientific). Surface rendering was performed by the volume tracer and color zone in UCSF ChimeraX.

### Manuscript preparation

The manuscript was written with the open and collaborative scientific writing package Manubot [29]. The source code and data for this manuscript are available at https://github.com/aseedb/synaptic_tomo_ms.

## Code availability

CryoVesNet is available on GitHub at https://github.com/Zuber-group/CryoVesNet

## Author contributions

BZ designed and supervised the study. RS and JR prepared samples, and acquired the data, reconstructed tomograms, and performed manual segmentation. AK, GW and BZ developed the CryoVesNet package. AK performed data analysis. JBS provided the SNAP-25 KO and mutant mice. RS and AK wrote the initial draft of the manuscript. AK, RS wrote the manuscript with contribution from all authors.

## Acknowledgments

This work was supported by the Swiss National Science Foundation (grant number 179520 to BZ), ERA-NET NEURON (NEURON-119 to BZ), and by the University of Bern Research Foundation (to Ioan Iacovache).

## Expanded View

### Expanded View Figures

**Figure EV1:**
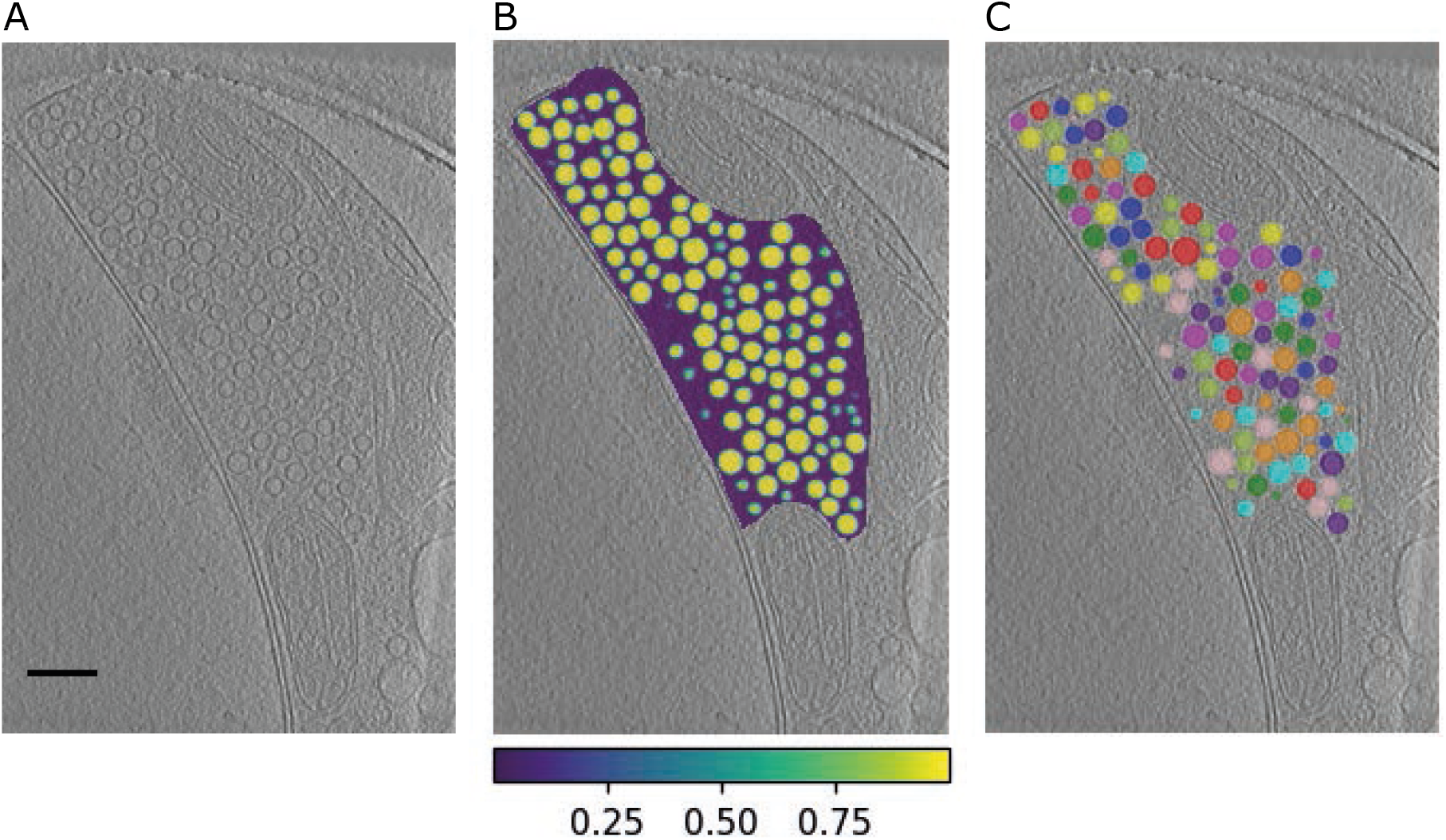
A 2D slice of an automatically segmented dataset. A) A section through a presynaptic terminal in a neuron tomogram. B) Predicted probability mask restricted to the segmentation region. Purple corresponds to a low SV probability and yellow to a high SV probability. C) Instance mask of the vesicles after post processing.

**Figure EV2:**
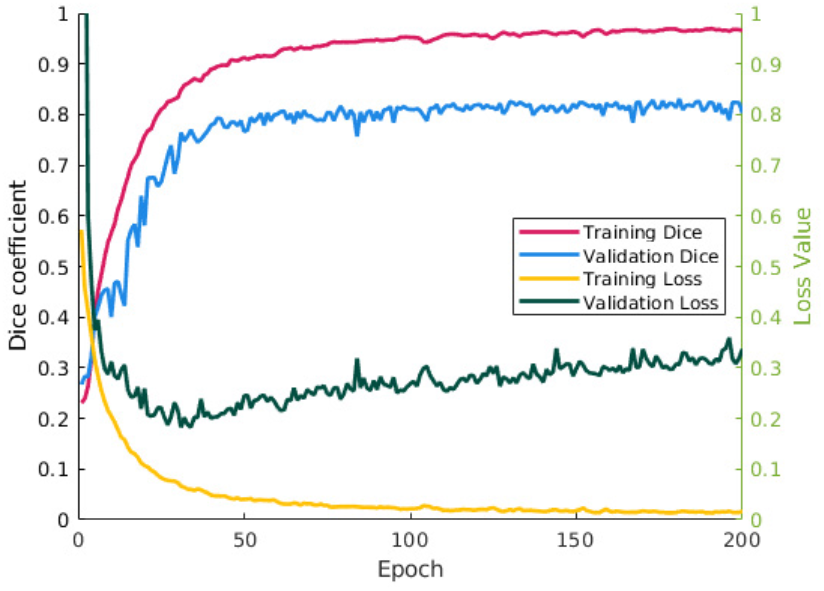
Dice coeficient and loss value for train and validation set.

### Expanded View Tables

**Table EV1:**
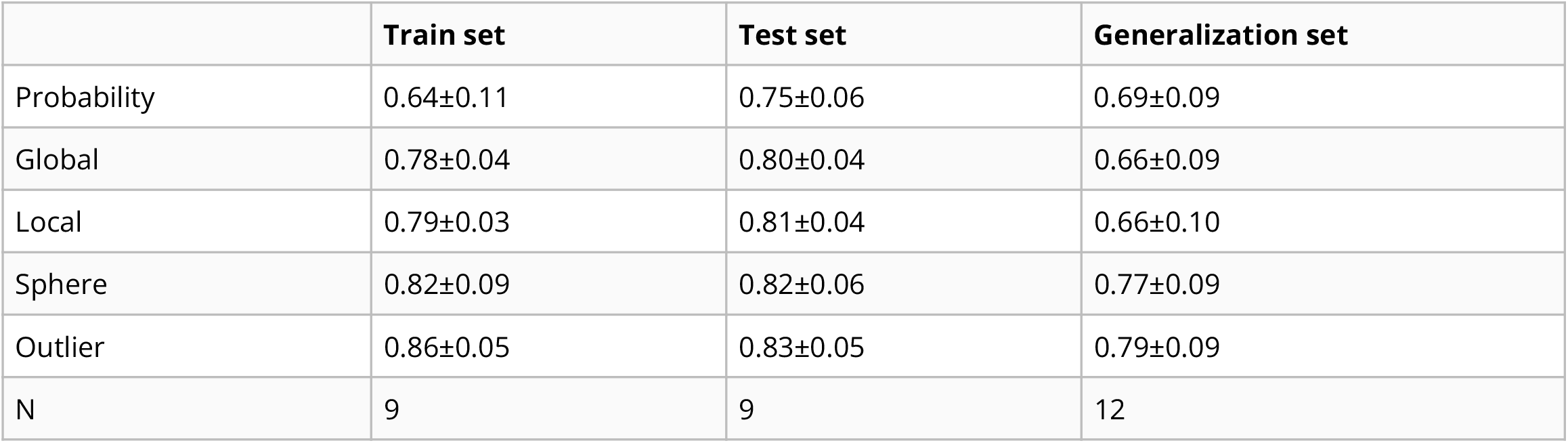
Dice statistical values. Mean ± standard deviation is shown at each step of the pipeline (Network Probability Mask, Global Threshold, Adaptative Localized Threshold, Sphere Radial Profile Refinement, and Outlier Removal).

**Table EV2:**
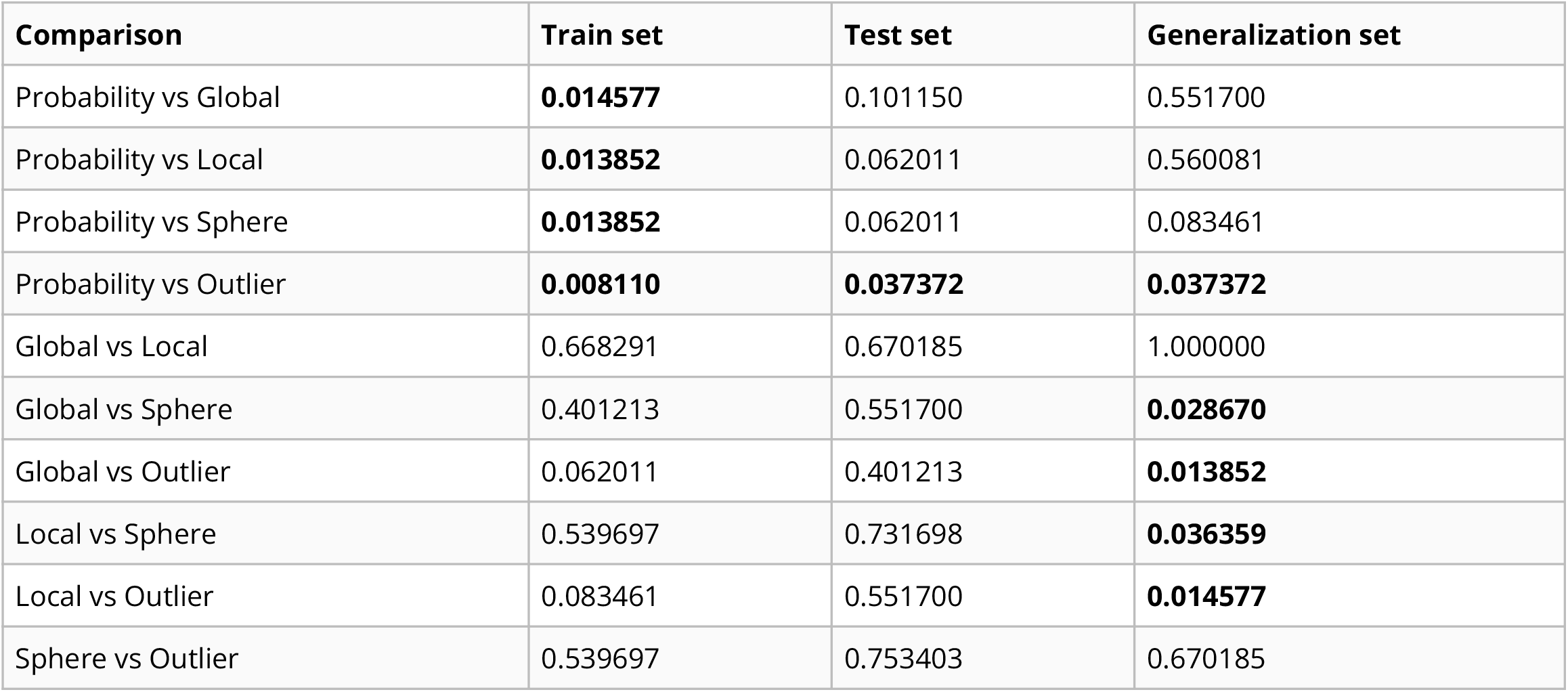
Corrected P-values of Dice values comparisons. Multiple, all-against-all ANOVA comparisons were performed with Benjamini-Hochberg correction on the Dice values summarized in EV1. Corrected P-values smaller than 0.05 are shown in bold.

